# A parametric finite element model of leg campaniform sensilla in *Drosophila* to study CS location and arrangement

**DOI:** 10.1101/2023.07.24.550300

**Authors:** Brian D. Saltin, Moritz Haustein, Ansgar Büschges, Alexander Blanke

**Affiliations:** Institute of Evolutionary Biology and Animal Ecology, University of Bonn, An der Immenburg 1, 53121 Bonn, Germany; Department of Animal Physiology, Institute of Zoology, University of Cologne, Zülpicher Str. 47b, 50674, Cologne, Germany

## Abstract

Campaniform sensilla (CS) are mechanosensors embedded within the cuticle of many insects at key locations such as nearby leg segment joints or halters. CS located at leg segments were found to respond to cuticle bending which can be induced by walking or jumping movements or by the underlying tensile forces of the muscles. For *Drosophila* it is unclear how CS location and material property variation influence stress levels within and around CS but this information is crucial to understand how flies might use CS input to adjust walking behaviour. Here, we designed a parametric model of the femoral CS field for *Drosophila* to allow for a systematic testing of the influence of CS location, orientation and material property variation on stress levels. The model consists of 7 changeable parameters per CS and 12 which can be changed for the CS field. Simulations of leg bending are in line with general beam bending theory: At the specific proximal CS field location nearby the trochantero-femoral leg joint, displacements are smaller than distal, while stresses are higher. When changing CS location towards more distal leg parts the situation changes towards more displacement and less stress. Changes in material property values for CS substructures or whole CS fields have a very low influence on stress or displacement magnitudes (regarding curve shape and amplitude) at the CS caps to which the nerve cells attach. Taken together, our simulation results indicate that for CS fields located at proximal leg parts, the displacements induced by other sources such as muscle tensile forces might be more relevant stimuli than the overall leg bending induced by typical locomotion scenarios. Future parametric finite element models should contain experimentally validated information on the anisotropic and viscoelastic properties of materials contained in this sensory system to further our understanding of CS activation patterns.

## 1 Introduction

The capabilities of insects to sense mechanical stress have been a field of interest for various biological disciplines ranging from comparative morphology all the way to robotics (Chapman, Duckrow and Moran, 1973; Field and Matheson, 1998; Zill, Büschges and Schmitz, 2011; Goldsmith, Szczecinski and Quinn, 2020).

Campaniform sensilla (CS) are a type of mechano-sensors which are located predominantly on the legs of many insects and the halteres of dipterans. CS consist of a convex cuticular cap which is often surrounded by a collar that connects to the cuticle (Thurm, 1964; Keil, 1997). Internally, the dendrite of a neuron connects to the cap(Moran, Chapman and Ellis, 1971; Spinola and Chapman, 1975; Keil, 1997) so that deformations or movements of the cap activate the neuron (Spinola and Chapman, 1975; Grünert and Gnatzy, 1987; Sun *et al*., 2020). This anatomy allows to monitor very small displacements of cuticle surrounding the CS and, consequently, allow to monitor forces acting on the exoskeleton (Pringle, 1938; Zill *et al*., 2014). CS can be arranged in fields, groups, or as single sensilla and they are usually found in close proximity to joints (Dinges *et al*., 2021). Load feedback from CS is known to influence activity in the motor system of walking insects with respect to control of striding phases and adaptation to unexpected perturbations. Currently there is no concept based on the properties of individual CS regarding their role in this task, although there are very precise descriptions of CS on legs in individual insects such as the fruit fly. In addition, measuring the responses of single CS is a major challenge that is currently methodologically achievable only for some larger animals (cockroach, locust, stick insect) (Hofmann and Bässler, 1986; Newland and Emptage, 1996; Akay *et al*., 2004; Zill *et al*., 2010, 2014; Zill, Büschges and Schmitz, 2011).

The present study has two main aims. First, we developed a virtual CS model with high details based on a range of imaging methods in order to allow for a variety of testing procedures regarding relevant input forces for CS. This CS was developed parametrically, i.e., the morphometrics of the model can be changed within certain ranges to allow for differently sized or shaped CS and also different CS numbers per field. Therefore, one can test many different CS configurations as well as different CS field sizes and locations on a given leg or body segment. Secondly, using this parametric model, we aimed to address how the location of a given CS field might influence its response curve during typical locomotion scenarios such as walking.

## 2 Material & Methods

We used left hind legs of male *Drosophila melanogaster* flies to investigate the femoral CS field which contains 11 individual CS in most flies observed. Based on ultrathin sections, transmission scanning electron microscopy (TEM), and scanning electron microscopy (SEM), a parametric Computer Aided Design (CAD) model was assembled in COMSOL (v 5.3.). We used 3D tracking of walking behaviour to infer the vector orientations of forces occurring during leg stance phases which were then used as input vectors for the finite element simulations. The details of each of these methodological steps are as follows:

### Fixation, Embedding & Sectioning

Animals were cooled down in a freezer, then transferred to a modified Karnovsky fixative (Karnovsky, 1964). Legs were dissected under Karnovsky fixative, followed by fixation for another 6h in the fridge. After storage in PBS buffer, samples were post fixed with Osmium (30min) followed by a graded dehydration with Acetone and subsequent infiltration with the embedding medium Epon substitute 812 (Sigma Aldrich, Germany). Samples were cured at 65° for 3 days. Sectioning was performed following the method of Blumer (2002), with a Reichert Jung Ultracut E using a histo diamond knife (DIATOME, Switzerland). Samples were cut as 350nm semi-thin sections to verify CS position and with 60-70 nm thin sections for TEM. Sections were contrasted with uranyl and lead acetate using a nanofilm surface analysis.

### TEM Observations & Geometry Extraction

A Zeiss EM10 A/B was used to collect images of the sections. In order to extract the geometry, 20 points along the morphology (Figure 2b) were each measured 10x times. All measurements were centred by substracting the lowest (Y-coordinate) and leftmost point (X-coordinate) from each measurement prior to averaging.

**Figure 1.**
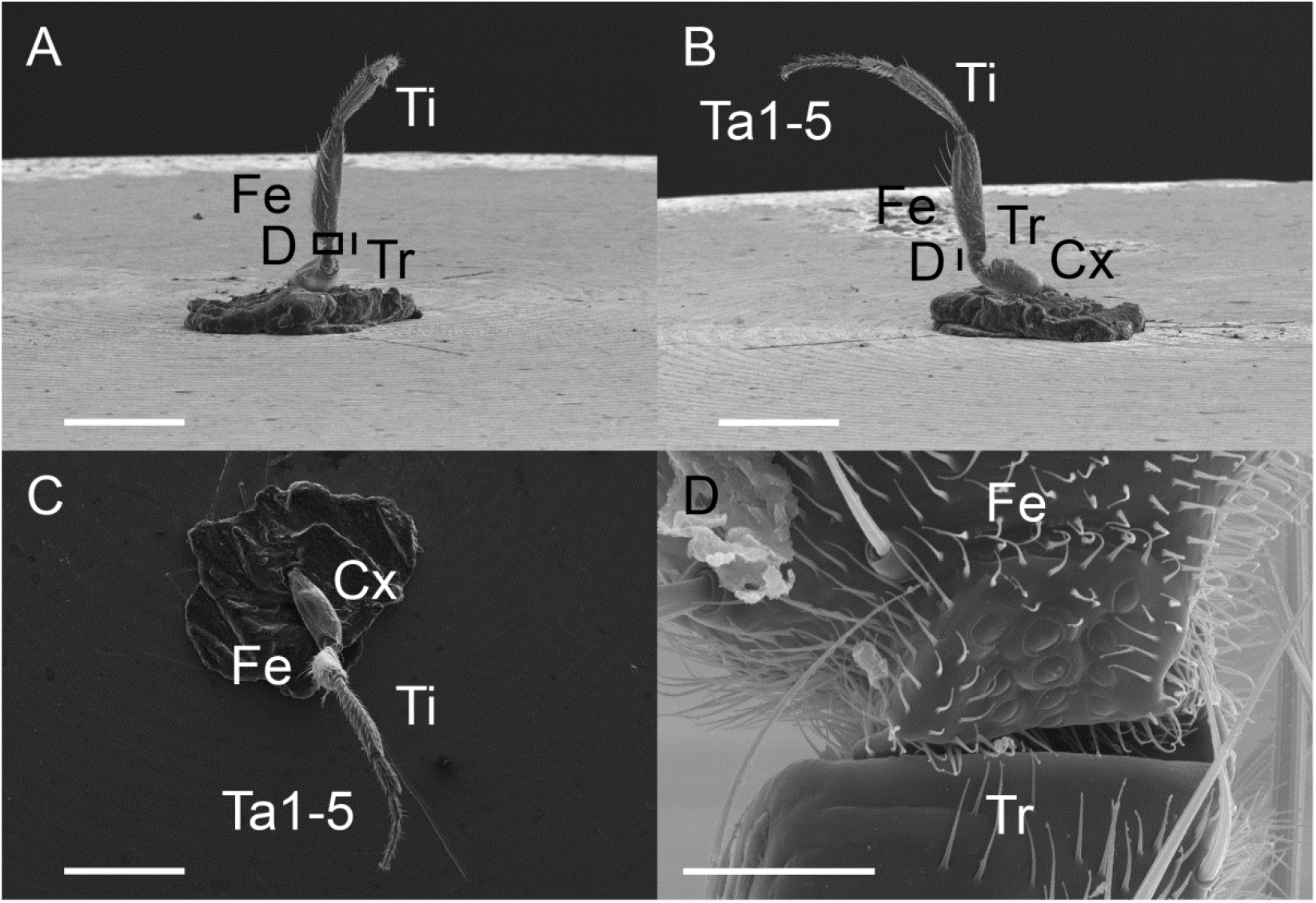
Exemplary location and arrangement of campaniform sensilla on the femur of *Drosophila melanogaster*. SEM Overview in 3D (A-C) and detail (D) of the shape and location of the femoral campaniform sensilla field in *Drosophila*. Scalebars are 500µm in A-C and 20µm in D. The location of D is marked in panel A & B. A and D are taken in the same orientation, B is rotated counter clockwise 90° around the Femur axis and C rotated 90° perpendicular to the femur axis towards the viewing plain in A. Ta1-5: tarsus one to five, Ti Tibia, Fe Femur, Tr Trochanter, Cx Coxa.

**Figure 2.**
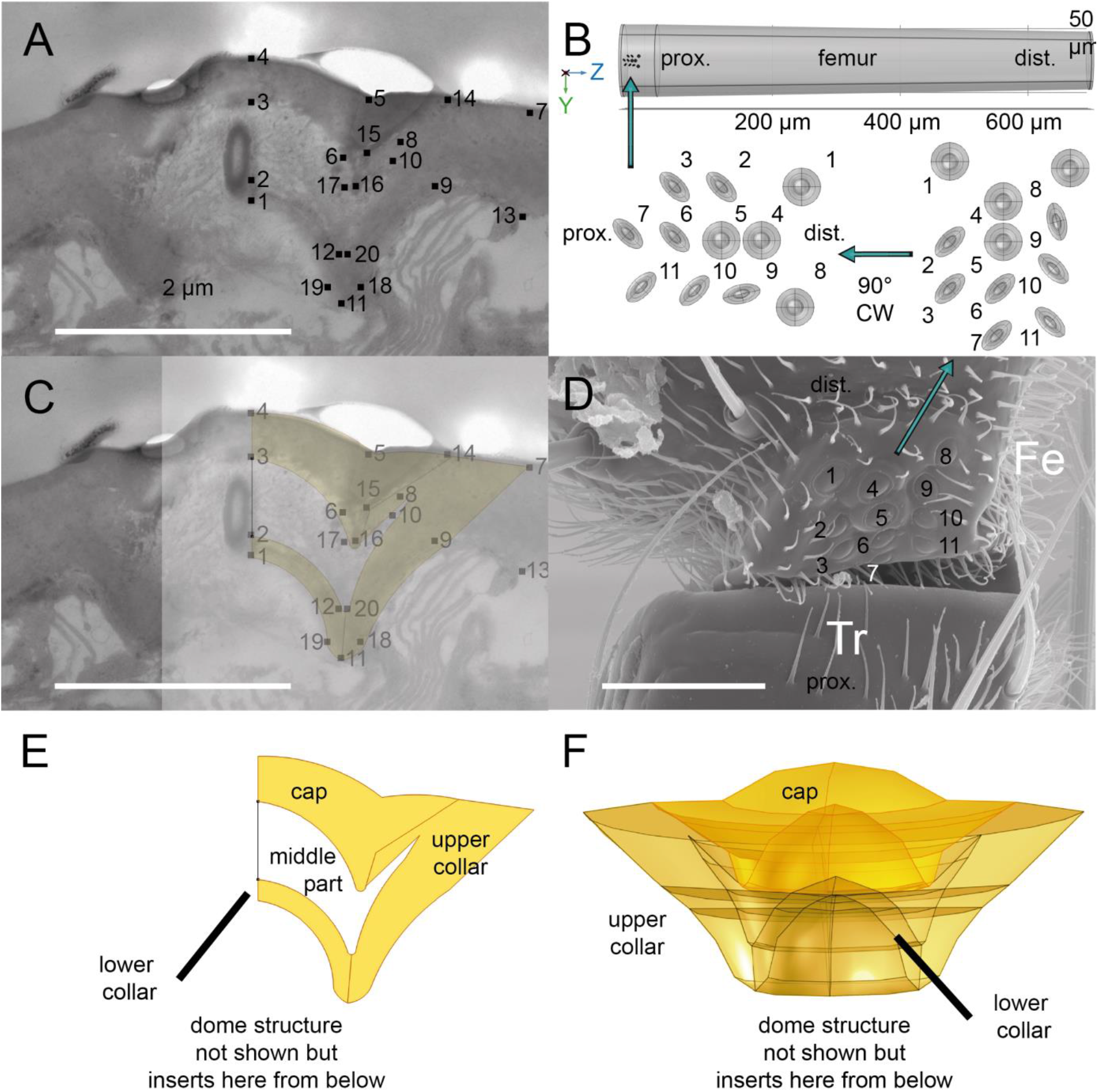
The location and arrangement of campaniform sensilla on the femur of *Drosophila melanogaster*. A) TEM image of a 60nm ultrathin section, through one CS showing the 20 points at which location data were extracted. B) Overview of a simplified femur model from proximal to distal, see also Figure 1. C) Projection of the 2D model generated in COMSOL onto the section. D) SEM overview, see Figure 1 for location, with numbering CS corresponding to the numbered CS models (multiplied from panel F) can been seen in panel B)) E) 2D model based on geometry extracted from the section. F) 3D rotation of the 2D model. Scale bar: A, C 2 µm and D 20µm.

### SEM Observations & Pattern Extraction

A Tescan Clara SEM was used to study the outer morphology of the femoral CS field at variable magnifications. Samples in 100% EthOH were dried at the critical point and sputter coated for 80 s with an acceleration voltage of 80 mV using a palladium target. In order to model length proportions of leg segments and substructures realistically in the parametric model, we performed a triangulation of the leg length using 3 orthogonal views during our SEM observations (Figure 1 A-C). The triangulation resulted in a length of 720,68 µm for the femur.

### CAD Model Development

Using the native CAD capabilities in COMSOL, the points extracted from TEM and SEM observations were entered in a so-called “work plain”, to reflect the rounded geometry (Figure 2). Adjacent points were connected with a circular arc. In a few cases (Figure 2) we used straight lines. Wherever possible, sharp edges were avoided as these are non-physiological, and also can cause singularities during the solving phase of FEA. CS were numbered from 1 (posterio-distal) to 11 (anterior-proximal) following Dinges (et al. 2022).

Individual sub-elements were handled independently to allow separate material allocation in all following steps. The CS model currently consists of a “cap”, “upper collar”, “lower collar”, a (reportedly “spongy” (Skordos *et al*., 2002)) “middle part” and the “dome” structure with the attached nerve.

To represent the whole CS field, the CAD CS model was multiplied and arranged to match the overall organisation of the femoral field and to match round and elliptic CS (Figure 2). Accessible animal independent parmeters can be found in Table 1. Triangulated leg segment lengthsI from SEM imaging were used to build a facsimile of a *Drosophila* femur and the field was warped to the leg circumference to achieve a perfect match of the outer circumference of each CS with the cuticle border. The current model supports one to twelve CS each with differing parameters (Table 1).

**Table 1:** Reference of the implemented parametric variables. Geometric variables used to generate CAD CS model in bold font. Changeable test variables in normal font. Already implemented variables not yet used in italics. Young’s modulus [YM] according to values from Skordos (2002).

| VariableName | Value | Numerical Variable Value | Description |
| --- | --- | --- | --- |
| Hblk1 | 30 | 30 | Height cylinder section as CS multiple Block1 (blk1) |
| XcMaster<br>_bend | 55 | 55 | Radius of cylinder_to_bend geometry around |
| FposCS1X to FposCS12X |  |  | X coordinate of CS1 to CS12 (of 11 in real case) as CS multiple 2D |
| FposCS1Y to FposCS12Y |  |  | Y coordinate of CS1 to CS12 (of 11 in real case) as CS multiple 2D |
| FposCS1Z to FposCS12Z |  |  | Z coordinate of CS1 to CS12 (of 11 in real case) as CS multiple 2D |
| FscaleCS1X to FscaleCS12X |  |  | scales height_usual do not touch. [range >0] 3D |
| FscaleCS1Y to FscaleCS12Y |  |  | Scales size of CS in Y if linked YZ plain [range >0] 3D |
| FscaleCS1Z to FscaleCS12Z | FscaleCS1Y to FscaleCS12Y | FscaleCS1Y to FscaleCS12Y | scale independently from YZ |
| FrotatedCS1 | 0[deg] | 0 rad | rotated CS1 |
| FrotatedCS2 | 0[deg] | 0 rad | rotated CS2 |
| FrotatedCS3 | 0[deg] | 0 rad | rotated CS3 |
| FrotatedCS4 | -77.5[deg] | -1.3526 rad | rotated CS4 |
| FrotatedCS5 | 0[deg] | 0 rad | rotated CS5 |
| FrotatedCS6 | 45[deg] | 0.7854 rad | rotated CS6 |
| FrotatedCS7 | -45[deg] | -0.7854 rad | rotated CS7 |
| FrotatedCS8 | 45[deg] | 0.7854 rad | rotated CS8 |
| FrotatedCS9 | 45[deg] | 0.7854 rad | rotated CS9 |
| FrotatedCS10 | -45[deg] | -0.7854 rad | rotated CS10 |
| FrotatedCS11 | 45[deg] | 0.7854 rad | rotated CS11 |
| FrotatedCS12 | 0[deg] | 0 rad | rotated CS12 |
| TestLoad | 8.963E-6/3 | 2.99E-06 | load magnitude [N] on leg (cylinder) femur-tibia joint |
| AxZ_rot_angle_between_XY | 90[deg] | 1.5708 rad | load direction in XY plain – azimuthal |
| AxY_rot_angle_between_ZX | 0[deg] | 0 rad | load direction in ZY plain – elevation |
| SpringBaseF [N/m] | 471,44 | 471,44 | sping base_femural-trochanter joint |
| Young's modulus [Pa] | 6E+09 | 6E+09 | hard cuticle structures |
| Young's modulus [Pa] | 4,8E+09 | 4,8E+09 | middle cuticle structures |
| Young's modulus [Pa] | 1,5E+09 | 1,5E+09 | basic cuticle structures |
| Poisson's ratio | 0.3 | 0.3 | Cuticle |
| Young's modulus [Pa] | 1,0E+04 | 1,0E+04 | soft tissue |
| Poisson's ratio | 0.485 | 0.485 | soft tissue |
| Joint_damping | 0,3 | 0,3 | *DampingCoefficient springbase_femoral_joint_not used* |
| vMles_sig | 1,76E+08 | 1,76E+08 | *VALUE_NEEDED_plasticity_initial yield stress_not used* |
| vMles_sig2 | 8,55E+07 | 8,55E+07 | *VALUE_NEEDED_plasticity_flow_stress_not used* |
| vMles_sat | 26 | 26 | *VALUE_NEEDED_plasticity_saturation coefficient_not used* |

### 3D Tracking of Walking Behaviour

To be able to determine tibial movements in relation to the femur during hind leg stance phases we performed all experiments with 3-8 days old adult Bolt-GAL4>UAS-CsChrimson *Drosophila melanogaster* flies (Bidaye *et al*., 2020). The animals were reared on a standard medium at 25°C and 65% humidity in a 12h:12h day:night cycle. Prior to experiments, the animals were transferred to new vials in which the food was soaked with 50 µL of a 100 mmol L^-1^ all-trans-Retinal solution (R2500; Sigma-Aldrich, RRID:SCR_008988) and were kept in the dark for at least three days.

For motion capture, tethered flies walked on a spherical treadmill (Berendes *et al*., 2016) and a red laser (658 nm) targeting the animal’s head was used for optogenetic stimulation of sustained forward walking using activation of CsChrimson. Leg movements were recorded by six synchronized high-speed cameras (acA1300-200um, Basler AG, Ahrensburg, Germany) equipped with 50 mm lenses (LM50JC1MS, Kowa Optical Products Co. Ltd. Nagoya, Japan). Cameras were positioned around the animal so that each body side was simultaneously captured by three cameras allowing the front, side, and rear of the animals to be recorded (Figure **3A**). Videos were recorded at 400 Hz and a resolution 896 by 540 pixels.

**Figure 3.**
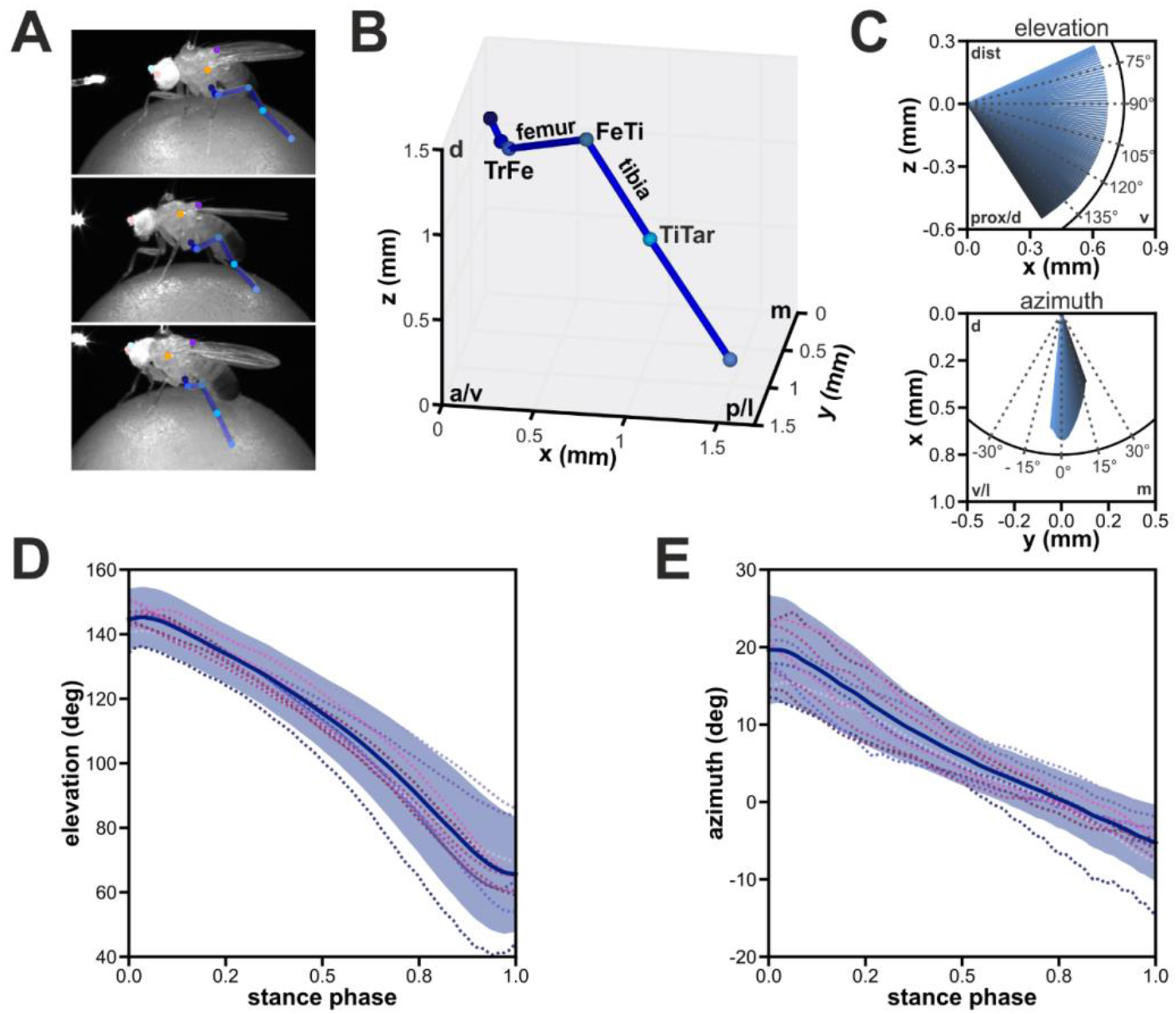
Tibial movements in relation to the femur during stance phases of the hind legs. (A) Representative images from a recorded walking sequence showing the front (upper panel), side (middle panel), and rear (lower panel) camera views of the left body side. For clarity, only the tracked positions of the hind leg are annotated. (B) 3D reconstruction of the posture of the left hind leg based on the tracked 2D positions of the joints and the tarsus tip in A. (C) Average movements of the tibia during the stance phase. The graphs display the mean orientation of the tibia (n = 224) in relation to femur. While the x-z plane, which corresponds to elevation angles, is depicted in the upper panel, the y-x plane, which corresponds to the azimuth, is displayed in the lower panel. The origin of both graphs is the femur-tibia joint (FeTi) – which is the position where, in the FE-simulation, forces transmitted by the tibia are acting upon. The time course is color-coded from begin (darker color) to end (brighter color) of the stance phase. (D+E) Time course of elevation (D) and azimuth (E) angles in relation to the stance phase. Blue solid line and area indicate mean and SD, while colored dotted lines represent the mean time courses of individual flies (N = 12). Abbreviations: a, anterior; p, posterior; v, ventral; d, dorsal; m, medial; l, lateral; dist, distal; prox, proximal.

For automated tracking of body parts in the videos, we used the DeepLabCut (DLC) toolbox (version: 2.2rc3; Mathis et al., 2018). We tracked six body parts on each leg: the thorax-coxa (ThCx) joint, coxa-trochanter (CxTr) joint, trochanter-femur (TrFe) joint, femur-tibia (FeTi) joint, tibia-tarsus (TiTar) joint and the tarsus tip (Tar). In addition, we tracked features on the fly’s body and head: the posterior scutellum apex on the thorax, the wing hinges, and the antennae. For this, we trained three independent ResNet-50 networks for the front, side, and rear camera groups, i.e. each network was able to track body parts in videos of both body sides from the same camera perspective. The used training sets contained 628, 655, 753 manually annotated images for the front, side, and rear networks, respectively. Resulting body feature positions were corrected manually as needed.

For 3D reconstruction of the positions of the body parts, we calibrated our camera setup with a custom-made checkerboard pattern (7 x 6 squares with size 399 µm x 399 µm per square) developed on a photographic slide (Gerstenberg Atelier für Visuelle Medien, Templin, Germany). For triangulation of 3D positions, a singular value decomposition algorithm was applied (Hartley and Zisserman, 2004; Günel *et al*., 2019). 3D positions of the tracked features were transformed to a body-centered coordinate system derived from the triangle formed by the wing hinges and the posterior scutellum apex.

To only analyse tibial movements of the hind legs during stance, the lift-off and touch-down events for each step were determined. By assuming that the distance between the tarsus tip and the center of the sphere on which the flies walked, must be approximately equal to the radius of the sphere (3 mm) during the stance phase, we performed a threshold operation (1.05-times the radius) to ascertain for each video frame whether a leg was in swing or stance phase. For this, the center position of the sphere was estimated by an optimization function which minimized the distance of tracked tarsus tips of all legs to the radius of the sphere. To ensure that tracked tarsus positions were not located inside the estimated sphere, a penalty factor of 100 was applied to tarsus tip distances smaller than the radius in the cost function.

To calculate the spherical coordinates, i.e. the elevation and azimuth angles, for the translation of tracked tibial movements to the FEA model, a local coordinate frame was established for each tracked hind leg posture in each analysed stance phase (Figure **3B+C**). In this context, elevation is correlated to tibial flexion and extension (i.e. 0° and 180° elevation angles correspond to fully-extended and fully-flexed, respectively), while the azimuth describes lateral motion of the tibia in relation to the femur. The origin was set to the position of the FeTi joint and the z-axis was derived from the orientation of the femur based on the positions of the TrFe and FeTi joints. The y-axis was derived from the normal of the femur-tibia plane. To be able to capture lateral movements of the tibia, i.e. the azimuth, a reference femur-tibia plane for each stance phase was used for all local coordinate frames of a stance phase in which the tibia was approximately orthogonal to the femur in the respective stance phase. To obtain right-handed coordinate frames, the x-axis was obtained by calculating the cross product of the vectors defining the z-axis and y-axis. For the right hind leg, the x-axis was inverted to allow the pooling of resulting spherical coordinates for both hind legs. Afterwards, the angles for elevation and azimuth of the tibia vector in each local coordinate frame were calculated (**Eqs. 1**).

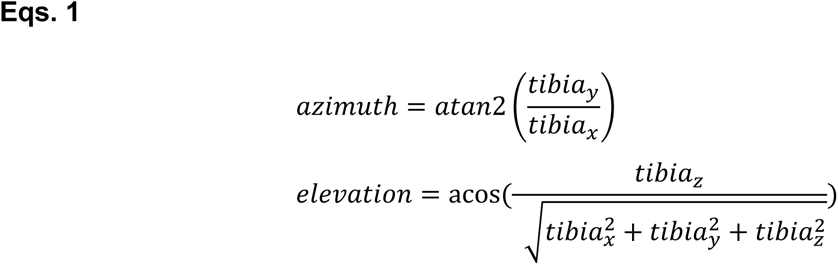

For calculating the mean time courses of elevation and azimuth across flies, the resulting angles were linearly interpolated to a sample size of 100. This resulted in normalized stance phases ranging from zero to one.

### Simulation Scenarios

Simulations were carried out with COMSOL 5.3 using the solid mechanics module. The results shown represent static analyses including only geometric nonlinearities. Given the lack of reliable experimental data for anisotropy and viscosity, a linear elastic material model was used. Although the simulations are static solutions, by solving the load distribution for different angular loadings, a full stepping cycle can be investigated. Therefore, we subjected each simulation to loading with 1/3 of 8.963 x 10^-6^ N which is the force resulting from the body weight of *Drosophila* (0.88 +/- 0.13mg). Forces were rotated in 7.5 deg steps along the Y-axis (reflecting “elevation”, i.e., dorso-ventral motion of the leg segment) and the Z-axis (azimuthal, i.e., anterior-dorsal motion) in order to simulate a complete walking cycle with a focus on the contact phase of the leg. Results were also verified with 1 degree steps for the stance phase’s angular range. The contact phase of the leg corresponded to 5.31° to 19.66° azimuthal and 144.58° to 65.57° elevation angle.

This general setup was used for several simulation scenarios: In **(A)** A cylinder representing the leg segment with no CS structure incorporated served as our baseline model. In **(B)** a cylinder with all CS substructure was modelled but all structures had the same material properties. In **(C)** we used a hollow cylinder with all CS substructure but additionally filled with a softer material to account for haemolymph and muscle material within the leg segment. Our **(D)** scenario contained the “full” model with different material values for each CS substructure.

In the following, we present the stress and displacement results for the central point on the “dome” encasing the neuron in order to show which stress and displacement levels the neuronal recording structures experience. Images were contrast and brightness optimized with Adobe Photoshop. Numerical data were exported from COMSOL to MATLAB (2023a) for further processing.

## 3 Results

### Angular Movement of the Tibia Segment of the Hind Leg During the Stance Phase

As in other walking insects, leg flexion and extension of the hind legs is mainly driven by tibial movements in *Drosophila*. During the stance phase the tibia is flexed at the beginning and extends until the end of the stance phase. These tibial movements corresponded to mean time course of elevation angles from our tracked motion capture data (Figure **3D**), starting at 144.58° ± 9.66° and followed by a progressive decrease to 65.57° ± 17.74° (n=244 stance phases from 12 tracked animals). Tibial movements showed also a lateral component in relation to the femur which was captured by the time course of the azimuth angles (Figure **3E**). Here, the mean azimuth angle was 19.66° ± 7.10° at the beginning of the stance phase and decreased steadily to −5.31° ± 5.00° during the stance phase.

### Finite Element Model validation

The overall curve progressions of displacements for all force vectors are the same for all simulation scenarios (Figure 4). Particularly noteworthy in this regard is that there is virtually no difference based on introduction of CS geometry elements provided they have the same material properties like the rest of the leg segment (<<1% difference in displacement magnitudes). The amplitude of the displacements is mainly influenced by filling of the hollow segment with a spongy material representing haemolymph and muscles. These results indicate a very low computational error due to element boundaries. The observed differences presented here are therefore due to changes in structural properties and/or material properties.

**Figure 4.**
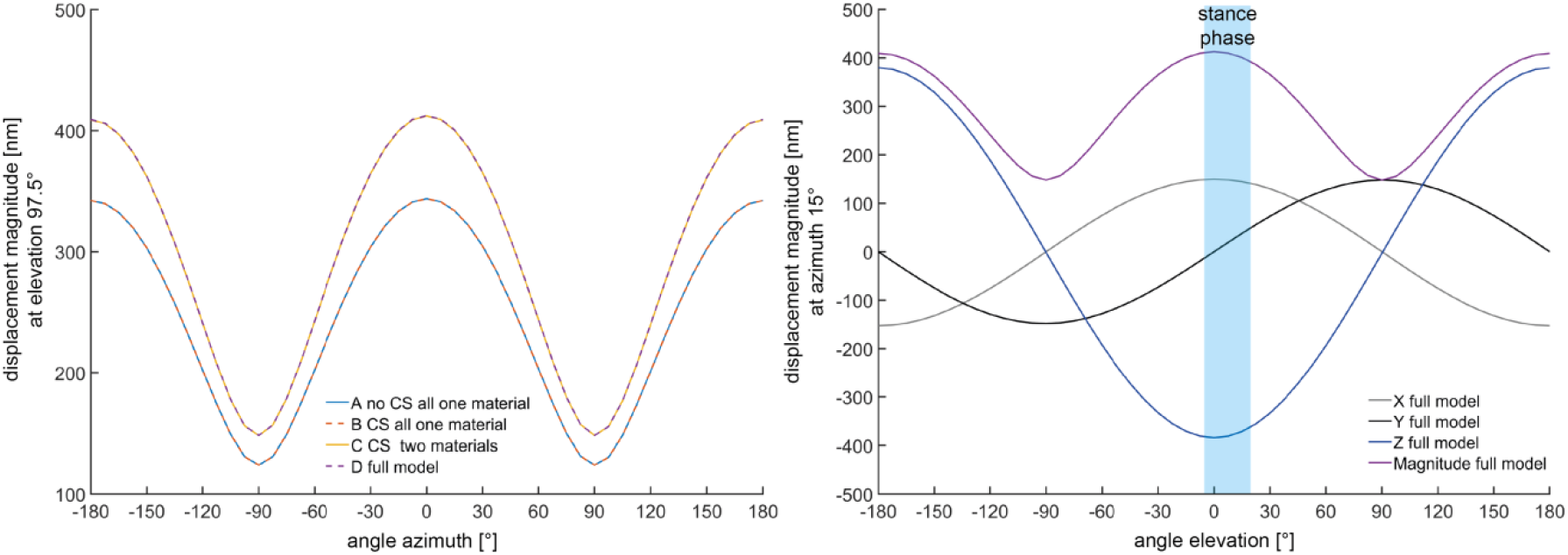
Displacement magnitudes for a full walking cycle for the 4 simulation scenarios A-D covering the full azimuthal range at 97.5 deg elevation. Note that the overall curve shape between “no CS on leg segment” (blue line) and the “full model” (purple line) are the same. The lower graph shows the absolute displacement decomposed into x-,y-, and z-components for the full model (scenario D). The overall pattern is the same for differing elevation angles (data not shown).

**Table 2:** Average percentage differences in total magnitude and individual X, Y, Z components of displacements for each of the three simulation scenarios B-D in relation to the baseline scenario A In round brackets: Percentage difference in total magnitude between (C-A) and (D-A).

|  | Scenario B in relation to A [%] | Scenario C in relation to A [%] | Scenario D in relation to A [%] |
| --- | --- | --- | --- |
| Magnitude of displacement | 0.00004303 | 16.40 | 16.45 (0.05) |
| X component | 0.00000717 | 20.76 | 21.19 (0.43) |
| Y component | 0.0000022 | 21.53 | 21.12 (0.41) |
| Z component | 0.00000223 | 17.92 | 17.91 (0.01) |

### Displacements at the CS Field During the Stance Phase

The femoral CS field is arranged in three columns (Figure 2, see also Dinges et al. 2022) with the following numbering used throughout this study: Anterior column with CS 1, 2 and 3, middle column with CS 4, 5, 6, and 7, and posterior column with CS 8, 9, 10, and 11 – each column is numbered low to high distal to proximal. The three CS columns display a clear phase lag in the azimuthal component regarding displacement (see Figure 5). CS rows, on the other hand, vary in amplitude, which is due to distance from the loading at the tibial-femoral joint.

**Figure 5.**
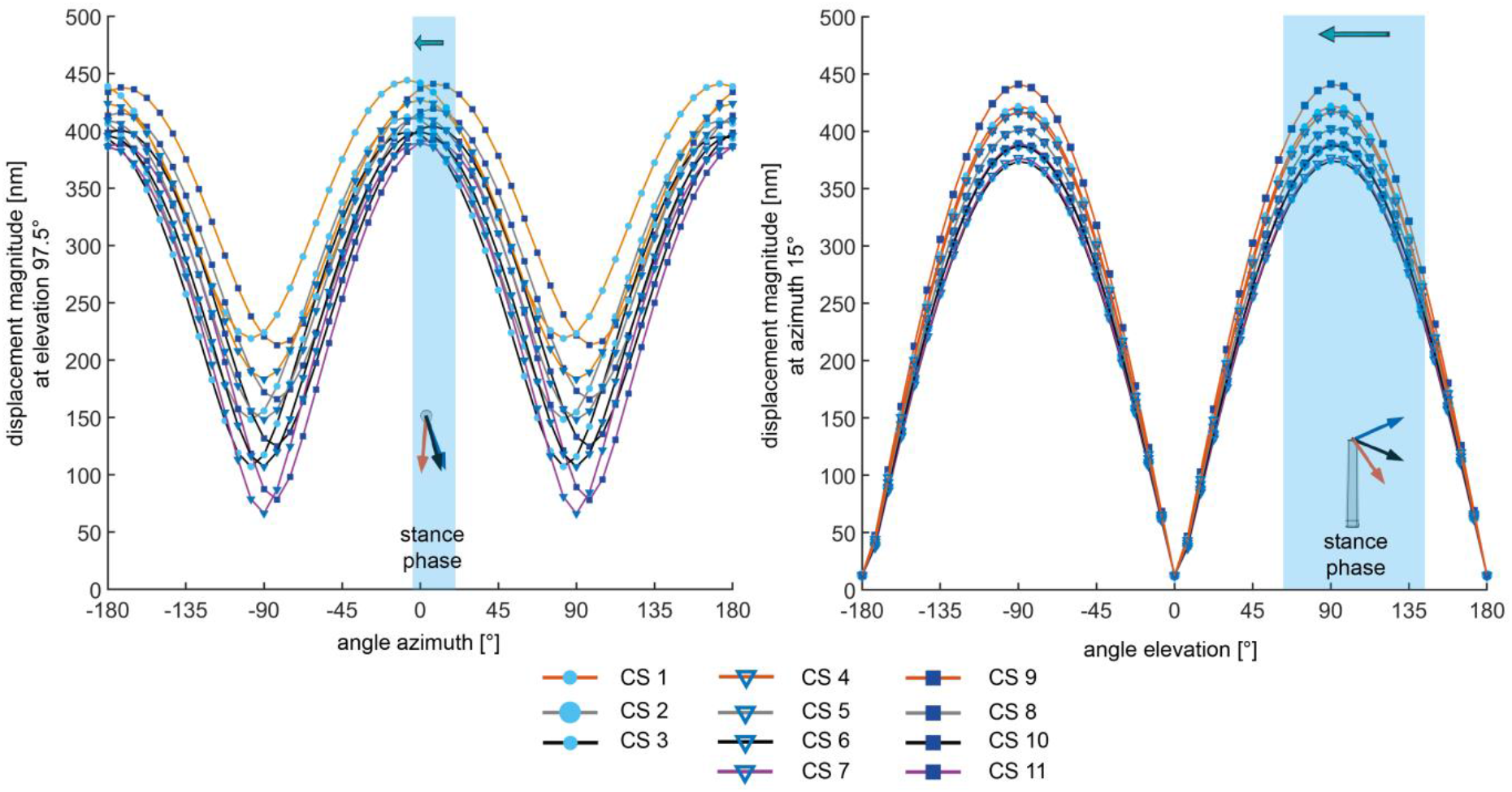
The left panel shows loading in azimuthal direction (X axis) vs displacement (Y axis) with each individual line representing an individual CS. The CS *columns* are colour-coded (blue boxes = anterior CS row, blue triangles = middle column, blue points = posterior column). *Rows* of CS are colour coded with line colour distal to proximal (orange, gray, black and purple). The right graph shows the same for elevation direction. The inset shows with a red arrow the start of motion and with a blue arrow the end of the motion. Black arrows indicate an elevation of 97.5° and azimuth 15°, respectively. The stance or contact phase of the leg is indicated by a white band the arrow at the top denotes the direction of motion in correspondence with figure 3.

During the stance phase of the leg (Figure 5), displacements both in elevation and azimuthal motion are maximal. This is true for all CS with the anterior-most CS column almost exceeding the angle range for contact. Surprisingly, the location of displacement maxima exemplary shown in Figure 5 is almost invariant with loading angle.

### Stress patterns at the CS Field During the Stance Phase

Comparing stress patterns for different force vector directions representative for walking phases (Figure 6) shows that amplitudes and the location of stress maxima do not vary substantially. This is true for relevant naturally occurring angular ranges in the femoral field for the principal most compressive stress (e3) and von Mises stress alike. The stress patterns also show that neither the shape (elliptical or circular) nor the orientation of single CS changes overall stress patterns for the natural simulation condition of a comparably distant force point relative to the location of the CS field. This counter intuitive result hints to the possibility that the effect of segment length and distance from loading needs further evaluation - an aspect studied in the following.

**Figure 6.**
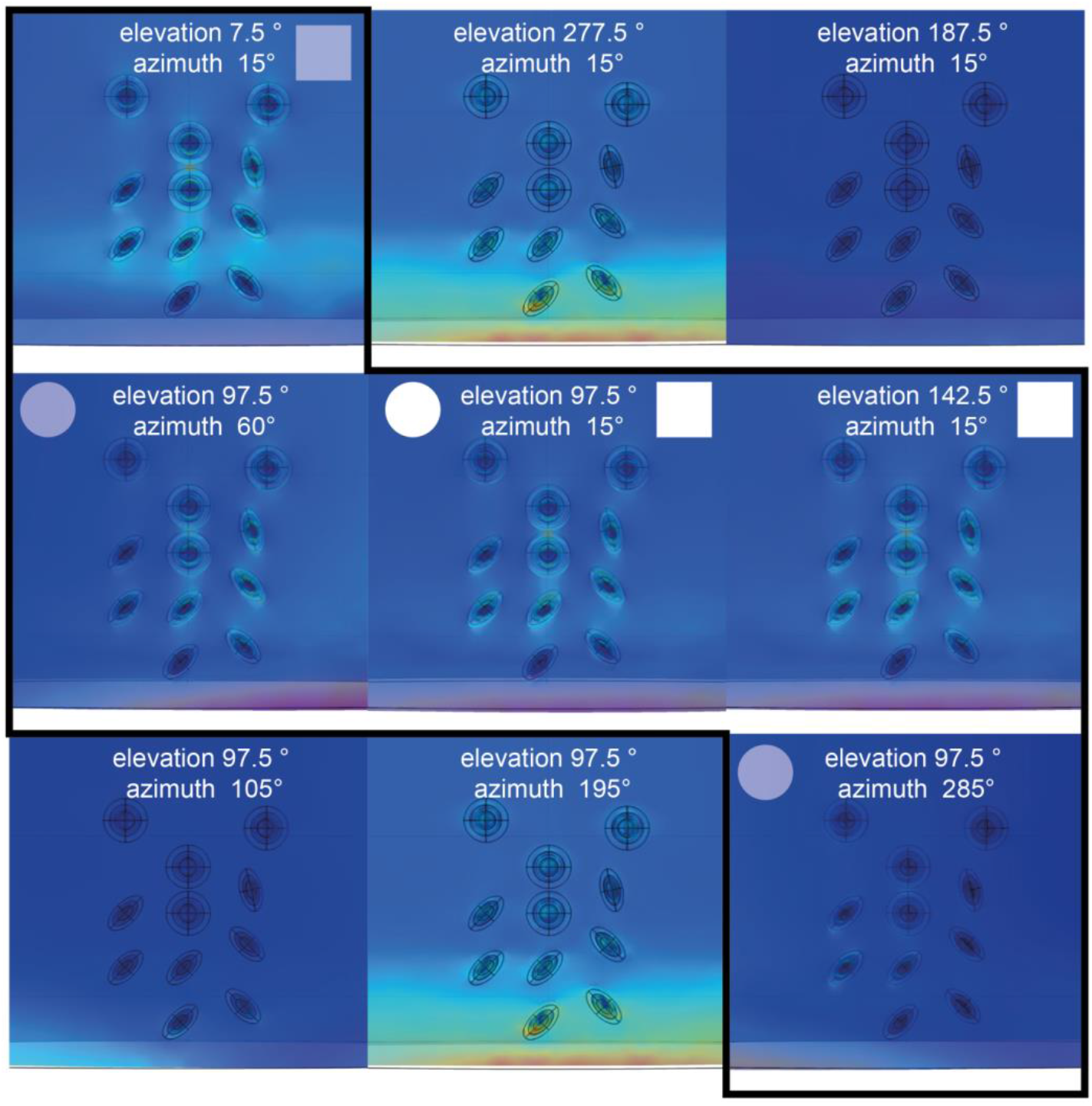
Qualitative overviews of principal compressive stresses for different combinations of elevation and azimuth angles relevant for walking in *Drosophila*. The central position represents the elevation (97.5°) and azimuth angle (15°) data for the graphs shown in Figure 4 & 5. The circle (for azimuth) and square (for elevation) in each upper corner denote if a given angle is part of the natural range (if full) or just outside (if shaded). The black line frames this information for both angles. Each angular view is set to its own range, in order to improve visibility of the qualitative pattern. Note that general stress distribution is invariant. Note that within in each angular combination, stress amplitudes are different which is due to the “auto range” in COMSOL for each combination, meaning comparisons of amplitude between individual CS in the field are possible, but not between different fields shown.

### Influence of segment length and point of loading relative to CS field location

To study the influence of leg segment length and relative position of the CS field, we used a simple cylinder (a simplified CS) incorporated almost the end of a lever arm (the leg segment) and varied the position of the cylinder along the lever arm from 5-95% (x-axis in Figure 7) relative to the point of force application (the femur-tibia joint). For each position of the cylinder along the lever arm we recorded the displacements for a typical gait cycle (y-axis in Figure 7). Results show that for spring suspended structures, from a certain distance on, the direction of force is inconsequential for the location of maximal displacement and minimal stress. The amplitude, however, varies with the force vector orientations dramatically. The simulations suggest that below 5-20% length of the femur, phases and amplitude have a strongly diminished effect. In this simple one armed lever structure, amplitude is influenced by distance. This confirms that the effect in the more complex model is physical and at natural leg segment length, forces transmitted from the step of the leg via the femoral-tibia joint in all likelihood at the real distance of over 650µm are not carrying directional information in the sense of amplitude and phase angles, as far as a single CS are concerned.

**Figure 7:**
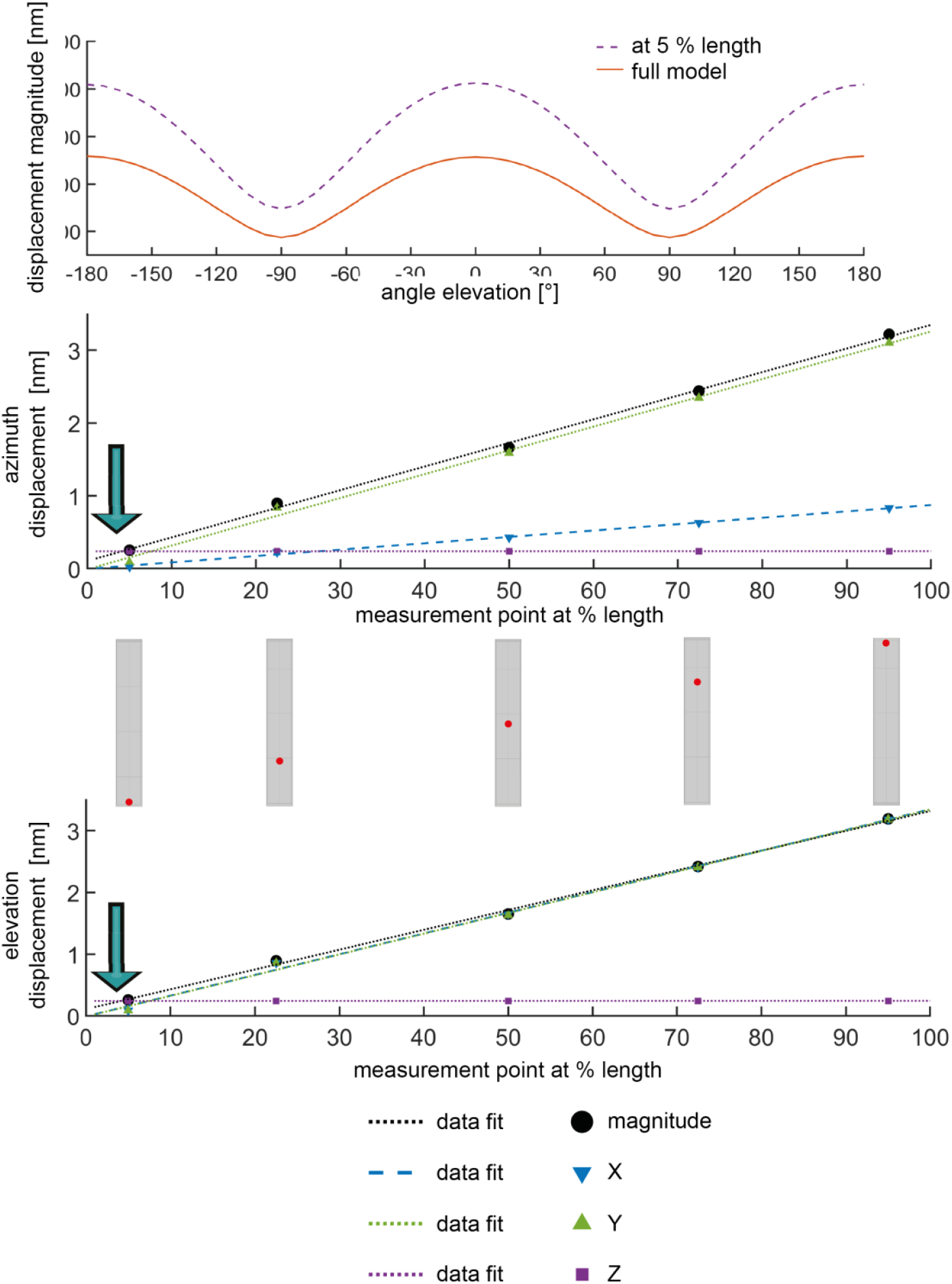
Displacements (azimuth and elevation) for a simplified model of a rectangular beam with a single CS (position denoted by red dot) at different locations along the beam. Points represent simulation results, lines the corresponding regressions. Displacement magnitude in black, x component of displacement in blue, y component in green, z component in purple. Note that the z-component is distance *independent*, while x- and y-components *decrease* depending on the CS location. Therefore, in the real femoral CS field in fruit flies, the z-component dominates displacement magnitudes since it is much larger than x & y. This is a likely explanation for the invariant compression patterns shown in figure 6. For the length of the femur (720 µm) this appears to be between 5-10% (36-72µm) in elevation and 5-20% in azimuth (>36µm). This is further distal in the leg than the real position of the femoral CS field in fruit flies.

## 4 Discussion

The present CAD/FE model unquestionably has its shortcomings – e.g. a geometry that needs refinement based on further TEM investigations. Also, we have not included damping effects and viscoelasticity, as well as anisotropy of the cuticle. The model also does not mimic perfectly the angular orientation of the CS, but given that the results indicate no drastic difference between elliptical and circular CS for the physiological load cases, this is not a major point of concern.

The model does, however, capture essential parts of the CS field’s mechanical behaviour and runs reliably within a sensible parameter range (Hepburn, 1976; Skordos *et al*., 2002; Vincent and Wegst, 2004; Dinges *et al*., 2022). Furthermore, the model allows to test the impact of variable morphologies of e.g. mutant lines and an investigation of CS configurations in other arthropods. The present results indicate a functional testing paradigm that in the future will allow comparing experimentally difficult to assess load cases with increased confidence. Most interestingly on a biological level, there are only very small variations of displacement maxima and only small effects of geometry of the CS, as well as direction of force, at the large natural distance of the CS to the femoral-tibia joint which is the point of force application in natural walking conditions. This has two major implications: First, a single CS, whether with a round or elliptical cap, has most likely no sensing capacity for different force directions at the femoral-tibia joint. The reaction forces occurring during walking are likely too small to be transferred into a leg bending that can be sensed by a femoral CS field far away from the point of force application.

Secondly, the above result hints to the possibility that this CS field (and possibly others in comparable positions) encode the cuticle displacements due to muscle tension. Muscle insertions and origins are located much nearer to the femoral CS field so that their activation might induce higher cuticle stresses and, consequently, also directional sensitivity. This dependence on muscle tension, if true, would be in line with the situation for the tibial campaniform sensilla in the cockroach which were found constitute a negative feedback system for the leg muscles (Zill and Moran, 1981). It was shown experimentally that these CS react to larger strains caused by resisted contractions of leg muscles (Zill and Moran, 1981).

For the stick insect middle leg it was shown that during forward walking, load signals from CS on the trochanter and femur initiate and maintain retractor and terminate protractor coxae activity (Akay *et al*., 2007). Furthermore, they initiate and maintain depressor and terminate levator trochanteris activity (Borgmann *et al*., 2011), and, finally, initiate flexor tibiae activity (Akay *et al*., 2001). Thereby, these CS assist the initiation and maintenance of the motor output required for stance movements. In contrast, CS signals mediating unloading of the leg support the transition to swing movement (reviewed in (Bidaye, Bockemühl and Büschges, 2018). Taken together, these earlier results for stick insects and the cockroach indicate that the CS investigated here might sense primarily loading (initiation of the stance phase) and unloading (initiation of the swing phase) via the cuticle displacements induced by muscle tension.

Another possibility is that the femoral CS field studied here only senses directionality when exposed to larger forces at the femoral-tibial joint. Such situations could occur during the initial jumping phase (Furuya et al. 2016) or during landing when reactions forces are ∼100x higher than during walking.

Although it remains unknown at which displacement/stress levels CS at the studied field and at other locations are activated, our model supports the view that not all CS might encode leg bending induced by locomotion reaction forces. Our future work will focus on the incorporation of more sophisticated material models (anisotropy and viscoelasticity) and possibly also neurophysiological experiments regarding activation thresholds for CS in *Drosophila*.

## Acknowledgements

We thank Sasha Zill,Nicholas S. Szczecinski, and Gesa Dinges for helpful comments and discussion. We also thank Thomas Bartolomaeus, Markus Koch and Tatjana Bartz for their support, insight and expertise regarding embedding and sectioning. MH is a member of the 233886668/GRK1960 (funded by DFG).

